## Supplementary Information for "Cryptic role of tetrathionate in the sulfur cycle: A study from Arabian Sea oxygen minimum zone sediments"

**Running Title:**	Tetrathionate metabolism in OMZ sediments

**Contents**

**Supplementary Tables**

**Supplementary Tables S1-S2.** Sediment-depths of SSK42/5 and 6 explored via duplicate metagenome sequencing and analysis of *in situ* microbial communities. Sequence accession numbers and basic statistics of the individual metagenome analyses are also given.

**Supplementary Table S3.** Number of thiosulfate to tetrathionate-conversion genes identified within the metagenome assemblies obtained for the individual sediment-samples of SSK42/5 and 6. Since this Supplementary Table is more than one page long it has been provided as an Excel sheet named Table S3, within the Excel Workbook named Supplementary Dataset.

**Supplementary Table S4.** Number of tetrathionate-oxidation genes identified within the metagenome assemblies obtained for the individual sediment-samples of SSK42/5 and 6. Since this Supplementary Table is more than one page long it has been provided as an Excel sheet named Table S4, within the Excel Workbook named Supplementary Dataset.

**Supplementary Table S5.** Number of tetrathionate-reduction genes identified within the metagenome assemblies obtained for the individual sediment-samples of SSK42/5 and 6. Since this Supplementary Table is more than one page long it has been provided as an Excel sheet named Table S5, within the Excel Workbook named Supplementary Dataset.

**Supplementary Tables S6-S7.** Pair-wise Pearson correlation and Spearman rank correlation coefficients calculated between the prevalence of two metabolic-types, or the prevalence of a particular metabolic-type and sediment-depth, in SSK42/5 and 6.

**Supplementary Tables S8-S9.** Relative abundance along SSK42/5 and 6, for genera whose member species/strains have been reported for tetrathionate formation during their oxidation of thiosulfate to sulfate/ reduction of sulfate to sulfide.

**Supplementary Tables S10-S11.** Relative abundance along SSK42/5 and 6, for genera whose member species/strains have been reported for tetrathionate oxidation.

**Supplementary Tables S12-S13.** Relative abundance along SSK42/5 and 6, for genera whose member species/strains have been reported for tetrathionate reduction.

**Supplementary Tables S14-S17.** Results of slurry incubation experiments with the sediment-samples of SSK42/5 and SSK42/6 in thiosulfate-containing chemolithoautotrophic medium.

**Supplementary Table S18.** Results of slurry incubation experiments with the sediment-samples of SSK42/5 and SSK42/6 in tetrathionate-containing chemolithoautotrophic medium.

**Supplementary Table S19.** Genes for thiosulfate to tetrathionate conversion, tetrathionate-oxidation, and manganese-oxidation, identified within the contigs assembled from the metatranscriptomic sequence data obtained from the 275 cmbsf sample of SSK42/6.Since this Supplementary Table is more than one page long it has been provided as an Excel sheet named Table S19, within the Excel Workbook named Supplementary Dataset.

**Supplementary Table S20-S21.** Results of slurry incubation experiments with the sediment-samples of SSK42/5 and SSK42/6 in tetrathionate-containing heterotrophic medium.

**Supplementary Table S22.** Number of manganese-oxidation genes identified within the metagenome assemblies obtained for the individual sediment-samples of SSK42/5 and 6. Since this Supplementary Table is more than one page long it has been provided as an Excel sheet named Supplementary Table 22, within the Excel Workbook named Supplementary Dataset.

**Supplementary Methods**

Details of the methods used in Metagenome (total community DNA) sequencing.

**Supplementary References**

**Supplementary Tables**

**Table S1.** Sediment-depths of SSK42/5 explored via duplicate metagenome sequencing and analysis^1^ of *in situ* microbial communities.

| **Sediment-depths explored**  **(in cmbsf)** | **BioSample accession number** | **Sample-fraction** | **Run accession number** | **Data volume (in number of bases)** | **Total number of reads generated** | **Number of reads remaining after quality filtering, i.e. no. of reads available for annotation** |
| --- | --- | --- | --- | --- | --- | --- |
| 0 | SAMN04442175 | 1st fraction | SRR3646127 | 1,278,221,540 | 7,298,095 | 4,100,142 |
|  |  | 2nd fraction | SRR3646128 | 568,340,941 | 3,248,036 | 1,620,835 |
| 15 | SAMN04442176 | 1st fraction | SRR3646129 | 427,280,628 | 2,610,964 | 1,290,674 |
|  |  | 2nd fraction | SRR3646130 | 616,882,407 | 3,448,925 | 1,695,465 |
| 45 | SAMN04442177 | 1st fraction | SRR3646131 | 387,655,304 | 2,334,713 | 1,421,504 |
|  |  | 2nd fraction | SRR3646132 | 691,720,029 | 3,803,790 | 2,650,180 |
| 60 | SAMN04442178 | 1st fraction | SRR3646144 | 552,498,532 | 3,231,713 | 2,029,107 |
|  |  | 2nd fraction | SRR3646145 | 584,425,231 | 3,467,226 | 1,928,912 |
| 90 | SAMN04442179 | 1st fraction | SRR3646147 | 705,088,273 | 3,962,953 | 2,285,687 |
|  |  | 2nd fraction | SRR3646148 | 800,036,330 | 4,418,977 | 2,467,261 |
| 120 | SAMN04442180 | 1st fraction | SRR3646150 | 672,531,698 | 3,800,279 | 2,184,195 |
|  |  | 2nd fraction | SRR3646151 | 717,367,452 | 3,958,612 | 2,334,398 |
| 140 | SAMN04442181 | 1st fraction | SRR3646152 | 628,130,699 | 3,472,496 | 1,943,757 |
|  |  | 2nd fraction | SRR3646153 | 697,272,445 | 3,917,430 | 2,575,262 |
| 160 | SAMN04442182 | 1st fraction | SRR3646155 | 795,198,681 | 4,328,182 | 3,271,077 |
|  |  | 2nd fraction | SRR3646156 | 355,890,886 | 1,959,373 | 1,298,002 |
| 190 | SAMN04442183 | 1st fraction | SRR3646157 | 840,500,627 | 4,707,531 | 3,282,497 |
|  |  | 2nd fraction | SRR3646158 | 413,935,050 | 2,330,298 | 1,499,602 |
| 220 | SAMN04442184 | 1st fraction | SRR3646160 | 768,834,448 | 4,232,096 | 2,744,086 |
|  |  | 2nd fraction | SRR3646161 | 640,147,053 | 3,552,054 | 2,434,097 |
| 260 | SAMN04442185 | 1st fraction | SRR3646162 | 828,237,512 | 4,418,273 | 3,960,096 |
|  |  | 2nd fraction | SRR3646163 | 606,632,123 | 3,375,631 | 2,560,268 |
| 295 | SAMN04442186 | 1st fraction | SRR3646164 | 1,147,354,334 | 5,987,264 | 4,209,798 |
|  |  | 2nd fraction | SRR3646165 | 557,439,969 | 3,088,957 | 2,137,865 |

**^1^** Sequence accession numbers and basic statistics of the individual metagenome analyses are also given.

**Table S2.** Sediment-depths of SSK42/6 explored via duplicate metagenome sequencing and analysis^1^ of *in situ* microbial communities.

| **Sediment-depths explored**  **(in cmbsf)** | **BioSample accession number** | **Sample-fraction** | **Run accession number** | **Data volume (in number of bases)** | **Total number of reads generated** | **Number of reads remaining after quality filtering, i.e. no. of reads available for annotation** |
| --- | --- | --- | --- | --- | --- | --- |
| 2 | SAMN04442187 | 1st fraction | SRR3570036 | 1,230,228,553 | 7,526,987 | 2,746,941 |
|  |  | 2nd fraction | SRR3570038 | 686,577,413 | 4,013,116 | 1,664,995 |
| 30 | SAMN04442189 | 1st fraction | SRR3577067 | 237,352,587 | 1,494,448 | 328,185 |
|  |  | 2nd fraction | SRR3577068 | 329,077,937 | 1,756,452 | 697,608 |
| 45 | SAMN04442190 | 1st fraction | SRR3577070 | 666,231,908 | 4,215,458 | 1,123,949 |
|  |  | 2nd fraction | SRR3577071 | 867,564,670 | 4,835,349 | 2,071,474 |
| 60 | SAMN04442191 | 1st fraction | SRR3577073 | 404,688,873 | 2,487,315 | 591,960 |
|  |  | 2nd fraction | SRR3577076 | 555,206,604 | 2,944,982 | 1,261,964 |
| 75 | SAMN04442192 | 1st fraction | SRR3577078 | 864,644,833 | 4,862,257 | 2,679,721 |
|  |  | 2nd fraction | SRR3577079 | 1,391,798,499 | 7,689,069 | 5,510,682 |
| 90 | SAMN04442193 | 1st fraction | SRR3577081 | 138,012,396 | 766,150 | 442,790 |
|  |  | 2nd fraction | SRR3577082 | 1,378,228,859 | 7,813,905 | 5,847,429 |
| 120 | SAMN04442195 | 1st fraction | SRR3577086 | 1,474,898,981 | 8,828,701 | 4,123,827 |
|  |  | 2nd fraction | SRR3577087 | 238,370,723 | 1,421,119 | 553,948 |
| 135 | SAMN04442196 | 1st fraction | SRR3577090 | 150,085,378 | 855,081 | 426,652 |
|  |  | 2nd fraction | SRR3577311 | 1,336,528,154 | 7,483,917 | 5,299,184 |
| 175 | SAMN04442199 | 1st fraction | SRR3577337 | 715,363,715 | 4,459,448 | 1,541,170 |
|  |  | 2nd fraction | SRR3577338 | 960,992,630 | 5,159,276 | 3,031,479 |
| 220 | SAMN04442202 | 1st fraction | SRR3577341 | 529,204,581 | 3,293,193 | 1,117,704 |
|  |  | 2nd fraction | SRR3577343 | 761,724,326 | 4,036,862 | 2,253,858 |
| 250 | SAMN04442204 | 1st fraction | SRR3577344 | 246,259,982 | 1,605,212 | 770,221 |
|  |  | 2nd fraction | SRR3577345 | 320,216,160 | 1,901,213 | 214,455 |
| 265 | SAMN04442205 | 1st fraction | SRR3577347 | 191,837,021 | 1,201,578 | 294,881 |
|  |  | 2nd fraction | SRR3577349 | 246,735,820 | 1,391,322 | 499,764 |
| 275 | SAMN04442207 | 1st fraction | SRR3577350 | 589,206,520 | 3,704,141 | 1,499,641 |
|  |  | 2nd fraction | SRR3577351 | 764,712,806 | 4,307,878 | 2,693,134 |

**^1^** Sequence accession numbers and basic statistics of the individual metagenome analyses are also given.

**Table S6.** Pair-wise Pearson correlation coefficient (CC denoted as *r*) and Spearman rank correlation coefficient (RCC denoted as *ρ*) calculated between the prevalence of two metabolic-types, or the prevalence of a particular metabolic-type and sediment-depth, in SSK42/5.

| **Pearson correlation coefficient** | | | | | | | |
| --- | --- | --- | --- | --- | --- | --- | --- |
|  | Sediment depth | Mn-depositing bacteria | Mn-oxidizing bacteria | Tetrathionate-reducing bacteria | Tetrathionate-forming bacteria | Tetrathionate-oxidizing bacteria | ***P* value** |
| Sediment depth |  | 0.00012566 | 0.0305 | 2.06E-04 | 4.62E-04 | 6.99E-05 |  |
| Mn-depositing bacteria | 0.8858 |  | 0.0498 | 9.69E-07 | 3.24E-06 | 5.48E-07 |  |
| Mn-oxidizing bacteria | 0.6229 | 0.5764 |  | 0.013 | 0.0532 | 0.0412 |  |
| Tetrathionate-reducing bacteria | 0.8734 | 0.9579 | 0.6901 |  | 1.24E-06 | 2.12E-08 |  |
| Tetrathionate-forming bacteria | 0.85 | 0.9462 | 0.5696 | 0.9557 |  | 4.76E-06 |  |
| Tetrathionate-oxidizing bacteria | 0.899 | 0.9625 | 0.5951 | 0.9805 | 0.9418 |  |  |
|  | ***r* value** | | | | | |  |
| **Spearman rank correlation coefficient** | | | | | | | |
|  | Sediment depth | Mn-depositing bacteria | Mn-oxidizing bacteria | Tetrathionate-reducing bacteria | Tetrathionate-forming bacteria | Tetrathionate-oxidizing bacteria | ***P* value** |
| Sediment depth |  | 0.0014 | 0 | 9.70E-04 | 3.09E-04 | 0 |  |
| Mn-depositing bacteria | 0.8322 |  | 0.0047 | 5.94E-06 | 5.94E-06 | 9.17E-05 |  |
| Mn-oxidizing bacteria | 0.9091 | 0.7762 |  | 0 | 7.72E-04 | 0.002 |  |
| Tetrathionate-reducing bacteria | 0.8462 | 0.8951 | 0.9371 |  | 9.17E-05 | 5.97E-04 |  |
| Tetrathionate-forming bacteria | 0.8741 | 0.8951 | 0.8531 | 0.8881 |  | 1.92E-04 |  |
| Tetrathionate-oxidizing bacteria | 0.9301 | 0.8881 | 0.8182 | 0.8601 | 0.8811 |  |  |
|  | ***r* value** | | | | | |  |

**Table S7**. Pair-wise Pearson correlation coefficient (CC denoted as *r*) and Spearman rank correlation coefficient (RCC denoted as *ρ*) calculated between the prevalence of two metabolic-types, or the prevalence of a particular metabolic-type and sediment-depth, in SSK42/6.

| **Pearson correlation coefficient** | | | | | | | |
| --- | --- | --- | --- | --- | --- | --- | --- |
|  | Sediment depth | Mn-depositing bacteria | Mn-oxidizing bacteria | Tetrathionate-reducing bacteria | Tetrathionate-forming bacteria | Tetrathionate-oxidizing bacteria | ***P* value** |
| Sediment depth |  | 5.97E-01 | 0.0327 | 1.63E-01 | 3.30E-03 | 1.48E-02 |  |
| Mn-depositing bacteria | 0.1618 |  | 0.0333 | 2.91E-06 | 3.89E-01 | 2.59E-01 |  |
| Mn-oxidizing bacteria | -0.593 | 0.5914 |  | 0.1486 | 0.2907 | 0.5945 |  |
| Tetrathionate-reducing bacteria | 0.4109 | 0.9344 | 0.4241 |  | 6.36E-02 | 4.68E-02 |  |
| Tetrathionate-forming bacteria | 0.7483 | 0.2612 | -0.3174 | 0.528 |  | 8.02E-02 |  |
| Tetrathionate-oxidizing bacteria | 0.6566 | 0.3381 | -0.1631 | 0.5594 | 0.5024 |  |  |
|  | ***r* value** | | | | | |  |
| **Spearman rank correlation coefficient** | | | | | | | |
|  | Sediment depth | Mn-depositing bacteria | Mn-oxidizing bacteria | Tetrathionate-reducing bacteria | Tetrathionate-forming bacteria | Tetrathionate-oxidizing bacteria | ***P* value** |
| Sediment depth |  | 0.2973 | 0.0673 | 1.10E-01 | 1.40E-03 | 0.0044 |  |
| Mn-depositing bacteria | 0.3132 |  | 0.1634 | 0.00E+00 | 3.60E-02 | 2.17E-01 |  |
| Mn-oxidizing bacteria | -0.528 | 0.4121 |  | 0.2973 | 4.59E-01 | 0.4704 |  |
| Tetrathionate-reducing bacteria | 0.467 | 0.967 | 0.3132 |  | 7.50E-03 | 6.73E-02 |  |
| Tetrathionate-forming bacteria | 0.8077 | 0.5934 | -0.2253 | 0.7198 |  | 4.40E-03 |  |
| Tetrathionate-oxidizing bacteria | 0.7527 | 0.3681 | -0.2198 | 0.5275 | 0.7527 |  |  |
|  | ***r* value** | | | | | |  |

**Table S8.** Relative abundance along SSK42/5, for genera whose member species/strains have been reported for tetrathionate-formation during their oxidation of thiosulfate to sulfate / reduction of sulfate to sulfide.

|  |  | **Sediment-depths (in cmbsf)** | | | | | | | | | | | |  |
| --- | --- | --- | --- | --- | --- | --- | --- | --- | --- | --- | --- | --- | --- | --- |
| **Genera** | **Tetrathionate-forming phenotype** | **0** | **15** | **45** | **60** | **90** | **120** | **140** | **160** | **190** | **220** | **260** | **295** | **Reference** |
|  |  | **Mean relative abundance (as** **percentage of total metagenomic reads)** | | | | | | | | | | | |  |
| *Acidithiobacillus* | (i) S_2_O_3_ → S_4_O_6_ → SO_4_  (ii) S_4_O_6_ → SO_4_ | 0.06 | 0.04 | 0.04 | 0.07 | 0.497 | 0.421 | 0.24 | 0.12 | 0.164 | 0.326 | 0.178 | 0.267 | (Hedrich and Johnson, 2013) |
| *Advenella* | (i) S_2_O_3_ → S_4_O_6_ → SO_4_  (ii) S_4_O_6_ → SO_4_ | 0 | 0 | 0 | 0 | 0 | 0 | 0 | 0 | 0.001 | 0 | 0 | 0 | (Ghosh et al., 2005) |
| *Desulfobulbus* | (i) SO_4_ → S_4_O_6_ → HS^-^ | 0.06 | 0.05 | 0.04 | 0.04 | 0.10 | 0.07 | 0.74 | 0.08 | 0.05 | 0.09 | 0.07 | 0.06 | (Sass et al., 1992) |
| *Desulfovibrio* | (i) SO_4_ → S_4_O_6_ → HS^-^ | 0.5 | 0.48 | 0.40 | 0.39 | 0.38 | 0.57 | 0.73 | 0.31 | 0.35 | 0.43 | 0.29 | 0.22 | (Sass et al., 1992) |
| *Halothiobacillus* | (i) S_2_O_3_ → S_4_O_6_ → SO_4_  (ii) S_4_O_6_ → SO_4_ | 0.03 | 0.02 | 0.05 | 0.04 | 0.22 | 0.26 | 0.19 | 0.256 | 0.275 | 0.267 | 0.291 | 0.36 | (Sievert et al., 2000) |
| *Pusillimonas* | (i) S_2_O_3_ → S_4_O_6_ → SO_4_  (ii) S_4_O_6_ → SO_4_ | 0.01 | 0.01 | 0.01 | 0.01 | 0.02 | 0.03 | 0.03 | 0.04 | 0.03 | 0.04 | 0.06 | 0.03 | This study |
| *Thiomicrospira* | (i) S_2_O_3_ → S_4_O_6_ → SO_4_  (ii) S_4_O_6_ → SO_4_ | 0.07 | 0.05 | 0.11 | 0.12 | 0.534 | 0.557 | 0.505 | 0.656 | 0.812 | 0.684 | 0.989 | 0.997 | (Watsuji et al., 2016) |

**Table S9.** Relative abundance along SSK42/6, for genera whose member species/strains have been reported for tetrathionate-formation during their oxidation of thiosulfate to sulfate / reduction of sulfate to sulfide.

|  |  | **Sediment-depths (in cmbsf)** | | | | | | | | | | | | | |  |
| --- | --- | --- | --- | --- | --- | --- | --- | --- | --- | --- | --- | --- | --- | --- | --- | --- |
| **Genera** | **Tetrathionate forming phenotype** | **2** | | **30** | **45** | **60** | **75** | **90** | **120** | **135** | **160** | **175** | **220** | **250** | **275** | **Reference** |
|  |  | **Mean relative abundance (as** **percentage of total metagenomic reads)** | | | | | | | | | | | | | |  |
| *Acidithiobacillus* | ((i) S_2_O_3_ → S_4_O_6_ → SO_4_  (ii) S_4_O_6_ → SO_4_ | 0.06 | 0.08 | | 0.11 | 0.09 | 0.15 | 0.18 | 0.19 | 0.20 | 0.28 | 0.23 | 0.34 | 0.16 | 0.12 | ( Hedrich and Johnson, 2013) |
| *Advenella* | ((i) S_2_O_3_ → S_4_O_6_ → SO_4_  (ii) S_4_O_6_ → SO_4_ | 0.00 | 0.01 | | 0.00 | 0.00 | 0.00 | 0.00 | 0.00 | 0.00 | 0.00 | 0.00 | 0.01 | 0.00 | 0.00 | ( Ghosh et al., 2005) |
| *Desulfobulbus* | (i) SO_4_ → S_4_O_6_ → HS^-^ | 0.06 | 0.04 | | 0.04 | 0.03 | 0.06 | 0.06 | 0.07 | 0.05 | 0.07 | 0.04 | 0.03 | 0.01 | 0.01 | ( Sass et al., 1992) |
| *Desulfovibrio* | (i) SO_4_ → S_4_O_6_ → HS^-^ | 0.48 | 0.25 | | 0.27 | 0.24 | 0.33 | 0.29 | 0.31 | 0.33 | 0.36 | 0.22 | 0.16 | 0.15 | 0.19 | ( Sass et al., 1992) |
| *Halothiobacillus* | (i) S_2_O_3_ → S_4_O_6_ → SO_4_  (ii) S_4_O_6_ → SO_4_ | 0.07 | 0.13 | | 0.26 | 0.18 | 0.27 | 0.37 | 0.35 | 0.39 | 0.92 | 0.52 | 0.55 | 5.98 | 2.58 | (Sievert et al., 2000) |
| *Pusillimonas* | (i) S_2_O_3_ → S_4_O_6_ → SO_4_  (ii) S_4_O_6_ → SO_4_ | 0.51 | 0.42 | | 0.39 | 0.37 | 0.03 | 0.03 | 0.56 | 0.03 | 0.00 | 0.48 | 1.05 | 0.43 | 0.48 | This study |
| *Thiomicrospira* | (i) S_2_O_3_ → S_4_O_6_ → SO_4_  (ii) S_4_O_6_ → SO_4_ | 0.21 | 0.35 | | 0.58 | 0.43 | 0.76 | 1.00 | 0.88 | 0.86 | 1.31 | 1.32 | 1.55 | 0.51 | 0.55 | ( Watsuji et al., 2016) |

**Table S10.** Relative abundance along SSK42/5, for genera whose member species/strains have been reported for tetrathionate-oxidation.

|  |  | **Sediment-depths (in cmbsf)** | | | | | | | | | | | |  |
| --- | --- | --- | --- | --- | --- | --- | --- | --- | --- | --- | --- | --- | --- | --- |
| **Genera** | **Tetrathionate oxidation phenotype** | **0** | **15** | **45** | **60** | **90** | **120** | **140** | **160** | **190** | **220** | **260** | **295** | **Reference** |
|  |  | **Mean relative abundance (as percentage of total metagenomic reads)** | | | | | | | | | | | |  |
| *Bosea* | S_4_O_6_ → SO_4_ | 0.00 | 0.00 | 0.01 | 0.00 | 0.00 | 0.00 | 0.00 | 0.01 | 0.00 | 0.01 | 0.00 | 0.00 | ( Das et al., 1996) |
| *Burkholderia* | S_4_O_6_ → SO_4_ | 0.00 | 0.71 | 0.64 | 0.87 | 1.15 | 1.01 | 0.78 | 1.02 | 0.78 | 0.94 | 0.81 | 1.01 | ( Wittke et al., 1997) |
| *Campylobacter* | S_4_O_6_ → SO_4_ | 0.06 | 0.06 | 0.05 | 0.06 | 0.04 | 0.06 | 0.09 | 0.05 | 0.19 | 0.07 | 0.09 | 0.04 | ( Voordouw et al., 1996) |
| *Hydrogenovibrio* | S_4_O_6_ → SO_4_ | 0.00 | 0.00 | 0.00 | 0.00 | 0.00 | 0.00 | 0.00 | 0.00 | 0.00 | 0.00 | 0.00 | 0.00 | ( NishiIhara et al., 1991) |
| *Pandoraea* | S_4_O_6_ → SO_4_ | 0.00 | 0.01 | 0.00 | 0.01 | 0.00 | 0.00 | 0.01 | 0.00 | 0.00 | 0.01 | 0.00 | 0.00 | ( Anandham et al., 2010) |
| *Paracoccus* | S_4_O_6_ → SO_4_ | 0.05 | 0.21 | 1.13 | 0.35 | 0.09 | 0.11 | 0.07 | 0.21 | 0.08 | 0.33 | 0.07 | 0.12 | ( Ghosh et al., 2006) |
| *Pseudaminobacter* | S_4_O_6_ → SO_4_ | 0.00 | 0.00 | 0.01 | 0.00 | 0.00 | 0.00 | 0.01 | 0.00 | 0.00 | 0.01 | 0.00 | 0.00 | ( Lahiri et al., 2006) |
| *Sulfurivirga* | S_4_O_6_ → SO_4_ | 0.00 | 0.00 | 0.00 | 0.01 | 0.00 | 0.00 | 0.00 | 0.00 | 0.01 | 0.00 | 0.01 | 0.00 | (Takai et al., 2006) |
| *Thiobacillus* | S_4_O_6_ → SO_4_ | 0.08 | 0.07 | 0.17 | 0.25 | 1.59 | 1.47 | 0.80 | 0.91 | 0.75 | 1.24 | 0.73 | 1.26 | (Wood and Kelly, 1991) |
| *Thiohalorhabdus* | S_4_O_6_ → SO_4_ | 0.00 | 0.00 | 0.01 | 0.00 | 0.00 | 0.00 | 0.01 | 0.00 | 0.00 | 0.01 | 0.00 | 0.00 | (Sorokin et al., 2008) |

**Table S11.** Relative abundance along SSK42/6, for genera whose member species/strains have been reported for tetrathionate-oxidation.

|  |  | **Sediment-depths (in cmbsf)** | | | | | | | | | | | | |  |
| --- | --- | --- | --- | --- | --- | --- | --- | --- | --- | --- | --- | --- | --- | --- | --- |
| **Genera** | **Tetrathionate oxidation phenotype** | **2** | **30** | **45** | **60** | **75** | **90** | **120** | **135** | **160** | **175** | **220** | **250** | **275** | **Reference** |
|  |  | **Mean relative abundance (as percentage of metagenomic reads)** | | | | | | | | | | | | |  |
| *Bosea* | S_4_O_6_ → SO_4_ | 0.00 | 0.01 | 0.00 | 0.01 | 0.00 | 0.00 | 0.00 | 0.01 | 0.00 | 0.00 | 0.00 | 0.00 | 0 | (Das et al., 1996) |
| *Burkholderia* | S_4_O_6_ → SO_4_ | 0.54 | 0.86 | 0.93 | 0.75 | 0.87 | 0.91 | 0.87 | 1.33 | 0.82 | 1.00 | 0.96 | 0.66 | 1.28 | (Wittke et al., 1997) |
| *Campylobacter* | S_4_O_6_ → SO_4_ | 0.06 | 0.05 | 0.05 | 0.04 | 0.06 | 0.07 | 0.08 | 0.11 | 0.05 | 0.04 | 0.02 | 0.04 | 0.03 | (Voordouw et al., 1996) |
| *Hydrogenovibrio* | S_4_O_6_ → SO_4_ | 0.00 | 0.00 | 0.00 | 0.00 | 0.00 | 0.00 | 0.00 | 0.00 | 0.03 | 0.02 | 0.04 | 0.00 | 0.00 | (NishiIhara et al., 1991) |
| *Pandoraea* | S_4_O_6_ → SO_4_ | 0.00 | 0.00 | 0.01 | 0.00 | 0.00 | 0.01 | 0.00 | 0.00 | 0.00 | 0.00 | 0.01 | 0.00 | 0.00 | (Anandham et al., 2010) |
| *Paracoccus* | S_4_O_6_ → SO_4_ | 0.18 | 0.53 | 0.53 | 0.60 | 0.28 | 0.16 | 0.18 | 0.15 | 0.20 | 0.09 | 0.11 | 0.71 | 0.35 | (Ghosh et al., 2006) |
| *Pseudaminobacter* | S_4_O_6_ → SO_4_ | 0.00 | 0.00 | 0.00 | 0.00 | 0.00 | 0.00 | 0.00 | 0.00 | 0.00 | 0.00 | 0.00 | 0.00 | 0.00 | (Lahiri et al., 2006) |
| *Sulfurivirga* | S_4_O_6_ → SO_4_ | 0.01 | 0.00 | 0.00 | 0.00 | 0.01 | 0.00 | 0.00 | 0.00 | 0.01 | 0.00 | 0.00 | 0.00 | 0.00 | (Takai et al., 2006) |
| *Thiobacillus* | S_4_O_6_ → SO_4_ | 0.22 | 0.34 | 0.66 | 0.47 | 0.91 | 1.19 | 1.02 | 0.92 | 0.98 | 1.27 | 1.26 | 0.31 | 0.34 | (Wood and Kelly, 1991) |
| *Thiohalorhabdus* | S_4_O_6_ → SO_4_ | 0.00 | 0.00 | 0.01 | 0.00 | 0.00 | 0.00 | 0.00 | 0.01 | 0.00 | 0.00 | 0.01 | 0.00 | 0.00 | (Sorokin et al., 2008) |

**Table S12.** Relative abundance along SSK42/5, for genera whose member species/strains have been reported for tetrathionate-reduction.

|  |  | **Sediment-depths (in cmbsf)** | | | | | | | | | | | |  |
| --- | --- | --- | --- | --- | --- | --- | --- | --- | --- | --- | --- | --- | --- | --- |
| **Genera** | **Tetrathionate reducing phenotype** | **0** | **15** | **45** | **60** | **90** | **120** | **140** | **160** | **190** | **220** | **260** | **295** | **Reference** |
|  |  | **Mean relative abundance (as percentage of total metagenomic reads)** | | | | | | | | | | | |  |
| *Alcaligenes* | S_4_O_6_ → S_2_O_3_ | 0.00 | 0.01 | 0.00 | 0.00 | 0.00 | 0.02 | 0.00 | 0.00 | 0.00 | 0.01 | 0.00 | 0.00 | (Barrett and Clark, 1987) |
| *Alteromonas* | S_4_O_6_ → S_2_O_3_ → H_2_S | 0.18 | 0.13 | 0.08 | 0.14 | 0.23 | 0.21 | 0.06 | 0.26 | 0.33 | 0.23 | 0.31 | 0.27 | (Barrett and Clark, 1987) |
| *Desulfotomaculum* | S_4_O_6_ → S_2_O_3_ → H_2_S | 0.32 | 0.28 | 0.28 | 0.15 | 0.20 | 0.23 | 0.40 | 0.08 | 0.16 | 0.21 | 0.08 | 0.05 | (Barrett and Clark, 1987) |
| *Desulfovibrio* | S_4_O_6_ → S_2_O_3_ | 0.49 | 0.47 | 0.40 | 0.40 | 0.38 | 0.58 | 0.73 | 0.32 | 0.36 | 0.20 | 0.29 | 0.23 | (Barrett and Clark, 1987) |
| *Edwardsiella* | S_4_O_6_ → S_2_O_3_ → H_2_S | 0.03 | 0.02 | 0.02 | 0.02 | 0.08 | 0.08 | 0.05 | 0.06 | 0.05 | 0.07 | 0.07 | 0.08 | (Barrett and Clark, 1987) |
| *Morganella* | S_4_O_6_ → S_2_O_3_ → H_2_S | 0.00 | 0.00 | 0.00 | 0.01 | 0.00 | 0.00 | 0.00 | 0.00 | 0.00 | 0.01 | 0.00 | 0.00 | (Barrett and Clark, 1987) |
| *Pasteurella* | S_4_O_6_ → S_2_O_3_ → H_2_S | 0.03 | 0.02 | 0.02 | 0.03 | 0.06 | 0.08 | 0.00 | 0.08 | 0.09 | 0.10 | 0.09 | 0.09 | (Barrett and Clark, 1987) |
| *Providencia* | S_4_O_6_ → S_2_O_3_ → H_2_S | 0.03 | 0.02 | 0.03 | 0.02 | 0.07 | 0.08 | 0.06 | 0.08 | 0.09 | 0.10 | 0.05 | 0.09 | (Barrett and Clark, 1987) |
| *Serratia* | S_4_O_6_ → S_2_O_3_ → H_2_S | 0.08 | 0.06 | 0.06 | 0.07 | 0.15 | 0.14 | 0.11 | 0.15 | 0.14 | 0.15 | 0.20 | 0.14 | (Barrett and Clark, 1987) |
| *Shewanella* | S_4_O_6_ → S_2_O_3_ | 1.21 | 0.83 | 0.70 | 1.26 | 1.87 | 1.84 | 1.69 | 2.09 | 3.42 | 2.00 | 3.04 | 3.50 | (Barrett and Clark, 1987) |

**Table S13.** Relative abundance along SSK42/6, for genera whose member species/strains have been reported for tetrathionate-reduction.

|  |  | **Sediment depths (in cmbsf)** | | | | | | | | | | | | |  |
| --- | --- | --- | --- | --- | --- | --- | --- | --- | --- | --- | --- | --- | --- | --- | --- |
| **Genera** | **Tetrathionate reduction phenotype** | **2** | **30** | **45** | **60** | **75** | **90** | **120** | **135** | **160** | **175** | **220** | **250** | **275** | **Reference** |
|  |  | **Mean relative abundance (as percentage of metagenomic reads)** | | | | | | | | | | | | |  |
| *Alcaligenes* | S_4_O_6_ → S_2_O_3_ | 0.00 | 0.01 | 0.00 | 0.00 | 0.44 | 0.00 | 0.00 | 0.00 | 0.01 | 0.00 | 0.01 | 0.02 | 0.01 | (Barrett and Clark, 1987) |
| *Alteromonas* | S_4_O_6_ → S_2_O_3_ → H_2_S | 0.23 | 0.19 | 0.22 | 0.19 | 0.28 | 0.31 | 0.27 | 0.36 | 0.23 | 0.30 | 0.23 | 0.17 | 0.23 | (Barrett and Clark, 1987) |
| *Desulfotomaculum* | S_4_O_6_ → S_2_O_3_ → H_2_S | 0.28 | 0.12 | 0.09 | 0.10 | 0.10 | 0.08 | 0.13 | 0.10 | 0.11 | 0.03 | 0.03 | 0.06 | 0.04 | (Barrett and Clark, 1987) |
| *Desulfovibrio* | S_4_O_6_ → S_2_O_3_ | 0.49 | 0.26 | 0.27 | 0.24 | 0.33 | 0.23 | 0.31 | 0.33 | 0.37 | 0.23 | 0.11 | 0.15 | 0.19 | (Barrett and Clark, 1987) |
| *Edwardsiella* | S_4_O_6_ → S_2_O_3_ → H_2_S | 0.03 | 0.05 | 0.06 | 0.04 | 0.06 | 0.08 | 0.07 | 0.09 | 0.08 | 0.09 | 0.13 | 0.02 | 0.04 | (Barrett and Clark, 1987) |
| *Morganella* | S_4_O_6_ → S_2_O_3_ → H_2_S | 0.00 | 0.00 | 0.00 | 0.00 | 0.01 | 0.00 | 0.00 | 0.00 | 0.01 | 0.01 | 0.00 | 0.00 | 0.00 | (Barrett and Clark, 1987) |
| *Pasteurella* | S_4_O_6_ → S_2_O_3_ → H_2_S | 0.04 | 0.06 | 0.07 | 0.05 | 0.08 | 0.12 | 0.10 | 0.10 | 0.07 | 0.09 | 0.10 | 0.04 | 0.04 | (Barrett and Clark, 1987) |
| *Providencia* | S_4_O_6_ → S_2_O_3_ → H_2_S | 0.04 | 0.06 | 0.07 | 0.06 | 0.09 | 0.11 | 0.10 | 0.12 | 0.08 | 0.09 | 0.09 | 0.03 | 0.06 | (Barrett and Clark, 1987) |
| *Serratia* | S_4_O_6_ → S_2_O_3_ → H_2_S | 0.08 | 0.10 | 0.13 | 0.12 | 0.13 | 0.16 | 0.16 | 0.20 | 0.18 | 0.20 | 0.22 | 0.10 | 0.17 | (Barrett and Clark, 1987) |
| *Shewanella* | S_4_O_6_ → S_2_O_3_ | 6.47 | 2.32 | 2.47 | 1.64 | 2.38 | 3.29 | 5.59 | 3.69 | 2.03 | 2.53 | 2.38 | 0.77 | 1.13 | (Barrett and Clark, 1987) |

**Table S14.** Rate of tetrathionate-formation in thiosulfate-containing chemolithotrophic ASWT medium by those sediment-samples of SSK42/5, which, in aerobic slurry culture experiments, oxidized thiosulfate only up to tetrathionate^1^.

| **Sediment-depths explored**  **(in cmbsf)** | **Rate of tetrathionate formation**  **(in μmol S day^-1^ g sediment^-1^)** |
| --- | --- |
| 0 | 17.72 |
| 15 | 6.65 |
| 90 | 8.42 |
| 160 | 6.45 |

^1^	These samples neither produced any sulfate from thiosulfate, nor further oxidized the tetrathionate produced from thiosulfate.

**Table S15.** Rate of formation, and subsequent oxidation (to sulfate), of tetrathionate during aerobic slurry incubation of various sediment-samples of SSK42/5 in thiosulfate-containing chemolithotrophic ASWT medium.

| **Sediment-depths explored**  **(in cmbsf)** | **Rate of tetrathionate formation from thiosulfate**  **(in μmol S day^-1^ g sediment^-1^)** | **Rate of oxidation of the tetrathionate formed from thiosulfate**  **(in μmol S day^-1^ g sediment^-1^)** |
| --- | --- | --- |
| 45 | 1.11 | 5.86 |
| 60 | 5.61 | 6.45 |
| 295 | 6.45 | 13.75 |

**Table S16.** Rate of tetrathionate-formation in thiosulfate-containing chemolithotrophic ASWT medium by those sediment-samples of SSK42/6 which, in aerobic slurry culture experiments, converted thiosulfate only to tetrathionate^1^.

| **Sediment-depths explored**  **(in cmbsf)** | **Rate of tetrathionate formation**  **(μmol S day^-1^ g sediment^-1^)** |
| --- | --- |
| 120 | 29.71 |
| 175 | 18.21 |
| 275 | 17.2 |

^1^	These samples neither produced any sulfate from thiosulfate, nor further oxidized the tetrathionate produced from thiosulfate.

**Table S17**. Rate of formation, and subsequent oxidation (to sulfate), of tetrathionate during aerobic slurry incubation of various sediment-samples of SSK42/6 in thiosulfate-containing chemolithotrophic ASWT medium.

| **Sediment-depths explored**  **(in cmbsf)** | **Rate of tetrathionate formation**  **(in μmol S day^-1^ g sediment^-1^)** | **Rate of tetrathionate**  **oxidation**  **(in μmol S day^-1^ g sediment^-1^)** |
| --- | --- | --- |
| 2 | 25.8 | 54 |
| 30 | 33.68 | 48 |
| 45 | 21.05 | 24 |

**Table S18**. Rate of oxidation (to sulfate) of tetrathionate during aerobic slurry incubation of various sediment-samples of SSK42/5 in tetrathionate-containing chemolithotrophic ASWTr medium.

| **Sediment-depths explored**  **(in cmbsf)** | **Rate of tetrathionate oxidation**  **(in μmol S day^-1^ g sediment^-1^)** |
| --- | --- |
| 0 | 23.5 |
| 15 | 4.75 |
| 45 | 3.35 |
| 90 | 4.9 |
| 120 | 5.71 |
| 160 | 2.5 |
| 295 | 6.4 |

**Table S20**. Rate of reduction of tetrathionate (to thiosulfate) during anaerobic slurry incubation of various sediment-samples of SSK42/5 in tetrathionate-containing heterotrophic RVTr medium.

| **Sediment-depths explored**  **(in cmbsf)** | **Rate of tetrathionate reduction**  **(in μmol S day^-1^ g sediment^-1^)** |
| --- | --- |
| 0 | 0.25 |
| 15 | 0.21 |
| 45 | 0.27 |
| 60 | 0.27 |
| 90 | 0.26 |
| 120 | 0.24 |
| 140 | 0.28 |
| 160 | 0.20 |
| 190 | 0.34 |
| 220 | 0.24 |
| 260 | 0.31 |
| 295 | 0.31 |

| **Sediment-depths explored**  **(in cmbsf)** | **Rate of tetrathionate reduction**  **(in μmol S day^-1^ g sediment^-1^)** |
| --- | --- |
| 2 | 0.35 |
| 30 | 0.28 |
| 45 | 0.5 |
| 60 | 0.58 |
| 75 | 0.49 |
| 90 | 0.86 |
| 120 | 1.2 |
| 135 | 1.28 |
| 175 | 1.35 |
| 220 | 1.5 |

**Table S21**. Rate of reduction of tetrathionate (to thiosulfate) during anaerobic slurry incubation of various sediment-samples of SSK42/6 in tetrathionate-containing heterotrophic RVTr medium.

**Supplementary Methods**

**Metagenome (total community DNA) sequencing**

Quality of metagenomic DNA samples was checked by electrophoresis and considered to be of high quality when no degradation signs were apparent. DNA quantity was determined using Qubit dsDNA HS Assay Kit (Thermo Fisher Scientific). 1 μg DNA from each sediment-sample was taken for deep shotgun sequencing by the Ion Proton platform using 200 bp read chemistry on a PI V2 Chip.

Libraries to be used for sequencing were constructed using the Ion Plus Fragment Library Kit (Thermo Fisher Scientific) and following the manufacturer’s Ion Plus gDNA and Amplicon Library Preparation User Guide. The Proton library was generated using 1 µg of genomic DNA which was fragmented to approximately 200 base pairs by the Covaris S2 system (Covaris, Inc., USA) and purified with 1.8x Agencourt Ampure XP Beads (Beckman Coulter, USA). Fragmentation was followed by end-repair, blunt-end ligation of the Ion Xpress Barcode and Ion P1 adaptors, and nick translation.

Post-ligation, size selection was done using E-Gel Size-Select 2% Agarose gels (Thermo Fisher Scientific) with the target size of 300 bp. Final PCR was performed using platinum PCR SuperMix High Fidelity and Library Amplification Primer Mix (custom product by Thermo Fisher Scientific), for 5 cycles of amplification. The resulting library was purified using AMPure XP reagent (1.2x; Beckman Coulter) and the concentration determined with Qubit dsDNA HS Assay Kit (Thermo Fisher Scientific); size distribution was done with Agilent 2100 Bioanalyzer high-sensitivity DNA kit (Agilent Technologies). Libraries were pooled in equimolar concentrations and used for template preparation.

Library templates were prepared for sequencing using OneTouch 2 protocols and reagents (Thermo Fisher Scientific). Library fragments were clonally amplified onto ion sphere particles (ISPs) through emulsion PCR and then enriched for template-positive ISPs. Proton emulsion PCR reactions utilized the Ion PI Template OT2 200 Kit v3 (Thermo Fisher Scientific). Following recovery, enrichment was completed by selectively binding the ISPs containing amplified library fragments to streptavidin coated magnetic beads, removing empty ISPs through washing steps, and denaturing the library strands to allow for collection of the template-positive ISPs. For all reactions, these steps were accomplished using the ES module of the Ion OneTouch 2. The selected ISPs were loaded on PI V2 Chip and sequencing was performed with the Ion PI 200 Sequencing Kit (Thermo Fisher Scientific) using the 500 flow (125 cycle) run format.
